# Asymmetric Distribution of Glucose Transporter mRNA Provides Growth Advantage

**DOI:** 10.1101/380279

**Authors:** Timo Stahl, Stefan Hümmer, Nikolaus Ehrenfeuchter, Geoffrey Fucile, Anne Spang

**Author notes:** Address of Correspondence: Anne Spang, Biozentrum, University of Basel, Klingelbergstrasse 70, CH-4056 Basel, Switzerland.

## Abstract

Asymmetric localization of mRNA is important for cell fate decisions in eukaryotes and provides the means for localized protein synthesis in a variety of cell types. Here we show that hexose transporter mRNAs are retained in the mother cell of *S. cerevisiae* until metaphase-anaphase transition (MAT) and then are released into the bud. The retained mRNA was translationally inactive but bound to ribosomes before MAT. Importantly, when cells were shifted from starvation to glucose-rich conditions, *HXT2* mRNA, but none of the other *HXT* mRNAs, was enriched in the bud after MAT. This enrichment was dependent on the Ras/cAMP/PKA pathway, the APC ortholog Kar9 and nuclear segregation into the bud. Competition experiments between strains that only expressed one hexose transporter at a time revealed that *HXT2* only cells grow faster than their counterparts when released from starvation. Therefore, asymmetric distribution of *HXT2* mRNA provides a growth advantage for young daughters, who are better prepared for nutritional changes in the environment. Our data provide evidence that asymmetric mRNA localization is an important factor in determining cellular fitness.

## Introduction

Cells have to respond to an ever-changing environment. This is true not only for single cell organisms like the yeast *Saccharomyces cerevisiae* but also for multicellular organisms, including humans. Variations in the availability of nutrients, in particular glucose, is one of the major challenges and cells have evolved a number of strategies to counteract glucose depletion. For example, under glucose-rich conditions insulin promotes the plasma membrane expression of the glucose transporter GLUT4 in adipocytes allowing glucose storage (Karnieli, Zarnowski et al., 1981, Martin, Millar et al., 2000). In contrast, another glucose transporter, GLUT1, is stabilized at the plasma membrane under energy stress conditions (Wu, Haynes et al., 2013). Alterations in glucose transporter expression have been observed in numerous diseases, including a variety of cancers. For example, GLUT1 and GLUT3 are overexpressed in solid tumors, and their expression levels are used as prognostic and predictive markers. Overexpression has been associated with poor survival (Barron, Bilan et al., 2016) as it may help to increase biomass production and tumor progression (Calvo, Figueroa et al., 2010, Yun, Rago et al., 2009).

The yeast *Saccharomyces cerevisiae* is sensitive to alterations of nutrient availability in the environment. Because of its inability to actively move towards a food source it has developed strategies to adapt quickly to local changes. Depending on the accessibility of glucose for example, yeast expresses a suitable set of its 17 hexose transporters to ensure an optimal growth pattern (Bisson, Fan et al., 2016). It has been well established that the transcription of the hexose transporters is modulated by glucose concentrations. In addition, the abundance of glucose transporters at the plasma membrane is regulated by endocytosis and subsequent degradation, if they are no longer needed (Hovsepian, Defenouillere et al., 2017, Llopis-Torregrosa, Ferri-Blazquez et al., 2016, O’Donnell, McCartney et al., 2015, Roy, Kim et al., 2014, Snowdon & van der Merwe, 2012). Even though these processes were initially studied mostly in yeast, they are conserved up to humans, indicating that *S. cerevisiae* is an excellent model organism for these types of studies.

Responses to changes in the environment can occur on both the transcriptional as well as the post-transcriptional level. Whereas our understanding of global transcriptional responses to environmental dynamics has vastly expanded from the deluge of next generation sequencing data, much less is known about post-transcriptional processes. This is partly due to the complexity of regulatory processes occurring at the levels of both RNA and protein. In the case of mRNA, a multitude of factors determines its stability, whether it is translated or stored, and how and where it is localized. All these mechanisms contribute to regulate protein expression and can be modulated in response to specific stresses (Wang, Schmich et al., 2018).

In this regard, subcellular localization of mRNA has been established as one important mechanism to control the timing and extent of protein expression (Parton, Davidson et al., 2014). During development and stem cell division for example, asymmetric localization of mRNA determines cell fate. Additionally, for neurotransmission, axonal mRNA localization and localized translation are essential. Yet, mRNA localization is not only limited to specialized cells. Localized mRNAs have been detected in a wide range of organisms including bacteria, yeast, plants and animals (Medioni, Mowry et al., 2012). Moreover, mRNA can be localized to a variety of intracellular organelles. In fact, there are thousands of mRNAs that exhibit specific subcellular localization patterns in mammals and Drosophila (Lecuyer, Yoshida et al., 2007, Mili, Moissoglu et al., 2008). It is generally assumed that mRNA localization correlates with regulated translation, suggesting that this highly conserved process is very efficient.

Here, we investigated the mRNA localization of the hexose transporter Hxt2. Hxt2 mRNA is asymmetrically localized during the cell-cycle, showing a strong retention in the mother cell until mitosis at which point equal portioning between mother and future daughter cell is achieved. The retained mRNA is presumably translational inactive during early phases of the cell-cycle and only becomes translated upon entry into mitosis. Importantly, refeeding of starved cells with glucose, shifted the equal distribution of *HXT2* mRNA to an enrichment in the bud. This process was dependent on the Ras/cAMP/PKA signaling pathway, spindle positioning and the nuclear pore components Nup2 and Mlp1/2, which were reported to be involved in nuclear pore complex inheritance. The asymmetric enrichment of *HXT2* mRNA during mitosis may contribute to a growth advantage of those daughters over cells that are unable to mount a similar response.

## Results

### Hxt2 protein is asymmetrically localized in yeast cells

We and others have used Hxt2 protein as a marker of the plasma membrane (Bagnat & Simons, 2002, Estrada, Muruganandam et al., 2015, Walther, Brickner et al., 2006, Zanolari, Rockenbauch et al., 2011). We observe, however, that in small and medium sized buds Hxt2 levels were always rather low but increased as the cell cycle progressed until at cytokinesis equal levels of Hxt2 were present in the mother and the daughter cells (Fig. 1A). These differences in protein levels might have been brought about by transcriptional and/or translational control. Indeed, several high throughput studies indicate that HXT2 mRNA expression occurs at restricted times in the cell cycle(Cho, Campbell et al., 1998, Pramila, Wu et al., 2006, Spellman, Sherlock et al., 1998). However, HXT2 mRNA was reported to be expressed at the mitosis to G1 boundary, an expression pattern that does not match well with the observed protein expression pattern. HXT2 mRNA is asymmetrically localized early and equally distributed late in the cell cycle.

**Figure 1:**
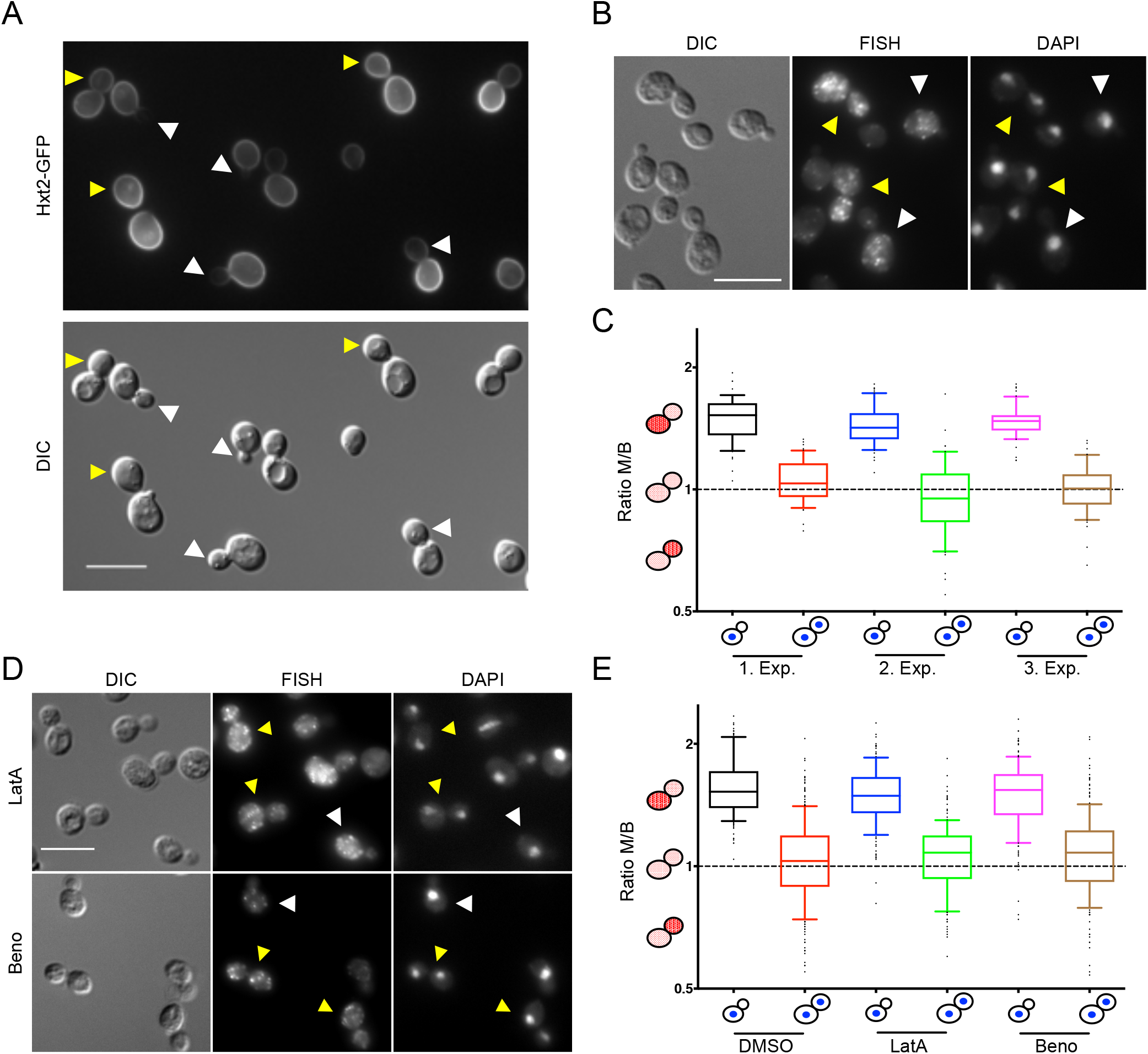
The localization of both HXT2 mRNA and Hxt2 protein is cell-cycle dependent. (**A**) The distribution of the GFP-tagged Hxt2 changes during the cell division. In small and medium budded cells (white arrowhead), Hxt2 is mainly present in the membrane of the mother cell. In large budded cells, Hxt2 is distributed over the entire plasma membrane (yellow arrowhead). (**B**) *HXT2* mRNA localization correlate with that of the protein. FISH with an HXT2-specific probe. Early in the cell cycle the mRNA is predominantly found in the mother cell while it is equally distributed after MAT (**C**) Quantification of (**B**). The fluorescence intensities from mother and bud were independently measured and the ratio of the signal from the mother over the signal from the bud was calculated. Values > 1 indicate a stronger signal in the mother, < 1 a stronger signal in the bud and around 1 equal distribution. Three independent experiments with 50 mother cells and 50 buds were analyzed. Boxes represent the interquartile range from the 25th to the 75th percentile with the median. Whiskers represent the 10th and 90th percentile, respectively. (**D**) Changes of HXT2 mRNA localization appear to be independent of the cytoskeleton. Cells were either treated for 15 min with 30 mg/ml latrunculin A or with 30 mg/ml benomyl. (**E**) Quantification of (**D**) of three in dependent experiments with at least 50 cells each. Scale bars in (A, B and D) correspond to 10 μm.

As pointed out above, mRNA localization is a potential mechanism to regulate protein expression. Therefore, we determined HXT2 mRNA cellular distribution by FISH (Fluorescence *In* Situ Hybridization) (Fig. 1B). In small and medium budded cells, HXT2 mRNA was restricted to the mother cell (Fig. 1B, white arrowheads), but in large budded cells (Fig. 1B, yellow arrowhead) HXT2 mRNA became equally distributed between mother and daughter cells. One explanation for this observation is that the mRNA distribution was connected to DNA segregation onto the two poles or in other words to the metaphase-anaphase transition (MAT). To investigate whether HXT2 mRNA localization is indeed correlated to cell cycle progression, we abrogated mitosis by treatment with nocodazole. Under these conditions, HXT2 mRNA remained restricted to the mother cell, suggesting a link between HXT2 mRNA localization and cell-cycle stage (Fig. S1A). For a more quantitative measure as readout from the FISH experiments, we determined the fluorescence intensity in the mother and the bud. The quotient of the mean fluorescence intensity of the mother cell over the bud/daughter cell reflects the relative mRNA distribution. A quotient of >1 indicates enrichment in the mother, and <1 in the bud (Fig. 1C). We scored cells with a bud and containing either one (before metaphase-anaphase transition (MAT)) or two nuclei (after MAT). We conclude that HXT2 mRNA localization changes over the cell cycle and that this change in localization is very robust and reproducible.

We wondered whether the HXT2 mRNA was transported to the bud through an active transport pathway, which would involve the cytoskeleton or through passive diffusion. We depolymerized actin cables with latrunculin A (LatA) (Fig. S1B) and microtubules (MTs) with benomyl (Fig. S1C), and assessed the localization of HXT2 mRNA. A benomyl concentration was chosen such that the number of cytoplasmic MTs was strongly reduced but cells could still undergo mitosis (Fig. S1C). However, neither mother cell retention nor release after MAT of HXT2 mRNA were affected when the cytoskeletal drugs were applied (Fig 1D and E). Our data suggest that under these conditions HXT2 mRNA diffuses into the bud after MAT.

### HXT2 mRNA localization in the bud is independent of de novo transcription

Even though Hxt2 mRNA was reported to be expressed at the M/G1 boundary, we cannot exclude that the change in distribution is due to transcriptional activity. To distinguish between mRNA movement in the cytoplasm from the mother to the bud and transcriptional increase of HXT2 mRNA concentration, we blocked transcription with 1,10-phenanthroline. Under these conditions the overall HXT2 mRNA signal was weaker, consistent with a decrease in transcriptional activity. Nevertheless, we still observed equal distribution of the mRNA after anaphase (Fig. S2A and B). Therefore, the change in HXT2 mRNA localization is largely independent of *de novo* mRNA synthesis. To corroborate our findings, we replaced the HXT2 promoter by the constitutive ADH and GPD promoters, which vary in their strength and led to a slight to moderate overexpression of HXT2 mRNA (Fig. S2C). Interestingly, under those conditions, HXT2 mRNA was still retained in the mother (Fig. S2D and E), indicating an active retention mechanism. In contrast, after mitosis, most of the HXT2 mRNA was still present in the mother cell, suggesting that the mRNA signal in the daughter is independent of increased mRNA levels. Taken together, these results suggest that transcription is not major contributor to HXT2 mRNA localization changes in logarithmically growing cells.

### HXT2 mRNA is associated with stalled ribosomes in the mother cell

Next, we asked whether these changes in mRNA localization were connected to translation. To this end, we treated cells with two translational inhibitors. Cycloheximide inhibits the elongation cycle during translation and therefore the mRNA is covered with stalled ribosomes (‘polysomes’), while verrucarin A is a translation initiation inhibitor, leading to accumulation of monosomes and ‘free’ RNA. In both cases, we observed a FISH signal reduction in the bud after MAT (Fig. 2A). In addition, we noticed that the overall FISH signal was stronger in cycloheximide-treated cells and somewhat weaker in the presence of verrucarin A, when compared to control (Fig. 2B), indicating that HXT2 mRNA is stabilized by binding to ribosomes. To test this hypothesis, we performed qPCR. Indeed, the HXT2 mRNA levels were slightly increased in the presence of cycloheximide (Fig. 2C). Conversely HXT2 mRNA levels were reduced when translation initiation was attenuated using the temperature-sensitive initiation factor *prt1-1* mutant (Fig. 2C). Our data imply that HXT2 mRNA requires ribosome association for protection from decay and suggest that HXT2 mRNA is ribosome-associated in the mother cell early in the cell-cycle. Since we detected Hxt2 protein only much later in the cell-cycle, we infer that the ribosomes on HXT2 mRNA would be stalled early in the cell cycle and that this block would be released at a later time point.

**Figure 2:**
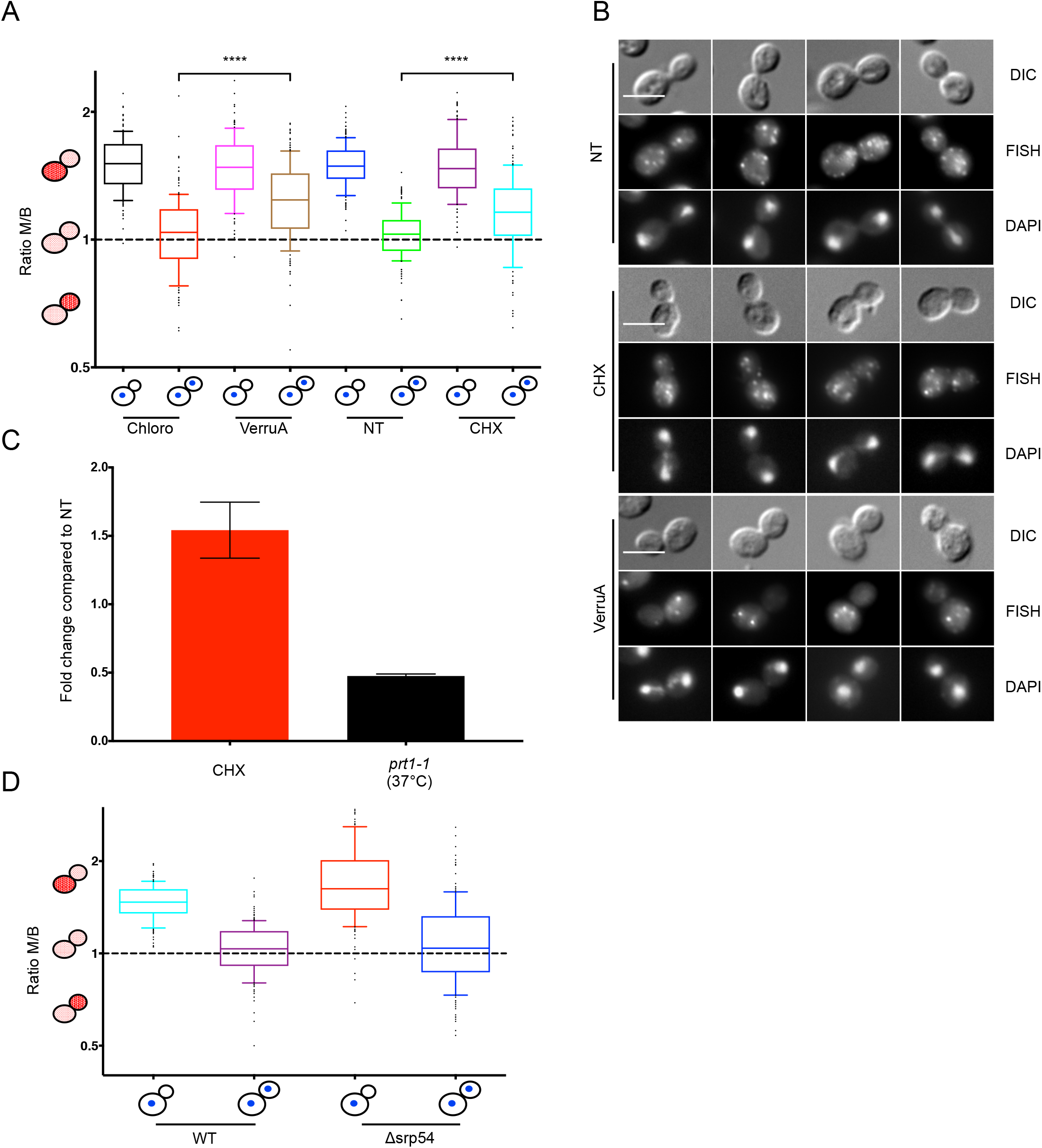
Active Translation is important for *HXT2* mRNA release to the bud. (**A** and **B**) Inhibiting translation by either applying Cycloheximide (CHX) or Verrucarin A (VerruA) leads to the retention of *HXT2* mRNA in the mother cell after MAT. FISH of CHX or VerruA-treated cells. Chloro: chloroform; solvent control for VerruA; NT: not treated; control for CHX (**C**) CHX treatment leads to increased transcript stability. Quantitative PCR of *HXT2* mRNA in CHX-treated cells in comparison to *prt1-1* mutant cells. (**D**) The localization of *HXT2* mRNA is independent of the Signal Recognition Particle (SRP). Deletion of *SRP54* does not affect the retention in the mother cell or the equal distribution of *HXT2* mRNA. Quantification of the FISH experiments in (**A**) and (**D**) was performed as in Fig. 1C. ****: P < 0.0001 in a twotailed, unpaired t-Test. Scale bars in (**B**) correspond to 5 μm.

### An HXT2 mRNA-ribosome complex is retained in the mother cell in an SRP-independent manner

Our data suggest that HXT2 mRNA is part of a translationally inactive ribosomal complex that is present in the mother cell before entry in mitosis. Since HXT2 mRNA is apparently synthesized in M/G1, one possible explanation for the mother cell localization is that HXT2 mRNA becomes ribosome-associated and translation is initiated in G1. The Hxt2 protein contains twelve transmembrane domains (Kasahara, Ishiguro et al., 2006) and hence must be co-translationally translocated into the ER. The transfer of a translating ribosome from the cytoplasm to the ER membrane is mediated by the signal recognition particle (SRP) (Hann & Walter, 1991). Thus, it is conceivable that during HXT2 mRNA translation, when the nascent polypeptide chain emerges from the ribosome, the HXT2 RNP is transferred to the ER by SRP, where the translation might be stalled. To test this possibility, we examined the HXT2 mRNA localization in a *Δsrp54* mutant, in which SRP function and translocation into the ER are strongly impaired (Hann & Walter, 1991). We did not observe any difference in the distribution of HXT2 mRNA between wild-type and *Δsrp54* cells (Fig. 2D), making it unlikely that SRP is actively involved in restricting HXT2 mRNA localization in the mother cell.

### Loss of the mRNA binding protein Scp160 causes enrichment of Hxt2 mRNA in the bud after mitosis

The results presented above imply that translation efficiency might be key to the retention of HXT2 mRNA in the mother cell. The polysome-associated mRNA binding protein Scp160 is ER-localized and is involved in translational efficiency (Hirschmann, Westendorf et al., 2014, Sezen, Seedorf et al., 2009, Weidner, Wang et al., 2014). Therefore, we tested whether *SCP160* deletion would affect HXT2 mRNA localization. While the localization was not altered early in the cell-cycle, HXT2 mRNA unexpectedly was enriched in the bud after MAT (Fig. 3A). *Δscp160* cells rapidly become polyploid (Wintersberger, Kuhne et al., 1995). To ensure that ploidy had no effect on the mRNA localization, we used a strain in which Scp160 expression is dependent on doxycycline (Hirschmann et al., 2014). Acute depletion of Scp160 also promoted HXT2 mRNA enrichment in the bud (Fig. 3B + Doxy, Fig S3A). Likewise deleting 2 or 4 of the C-terminal KH domains of Scp160 (Baum, Bittins et al., 2004) was sufficient for HXT2 mRNA bud enrichment (Fig. 3B). Thus, loss of Scp160 function promotes enrichment of HXT2 mRNA in the bud after anaphase. Yet, Scp160 does not appear to play a role in the initial retention of HXT2 mRNA in the mother cell.

**Figure 3:**
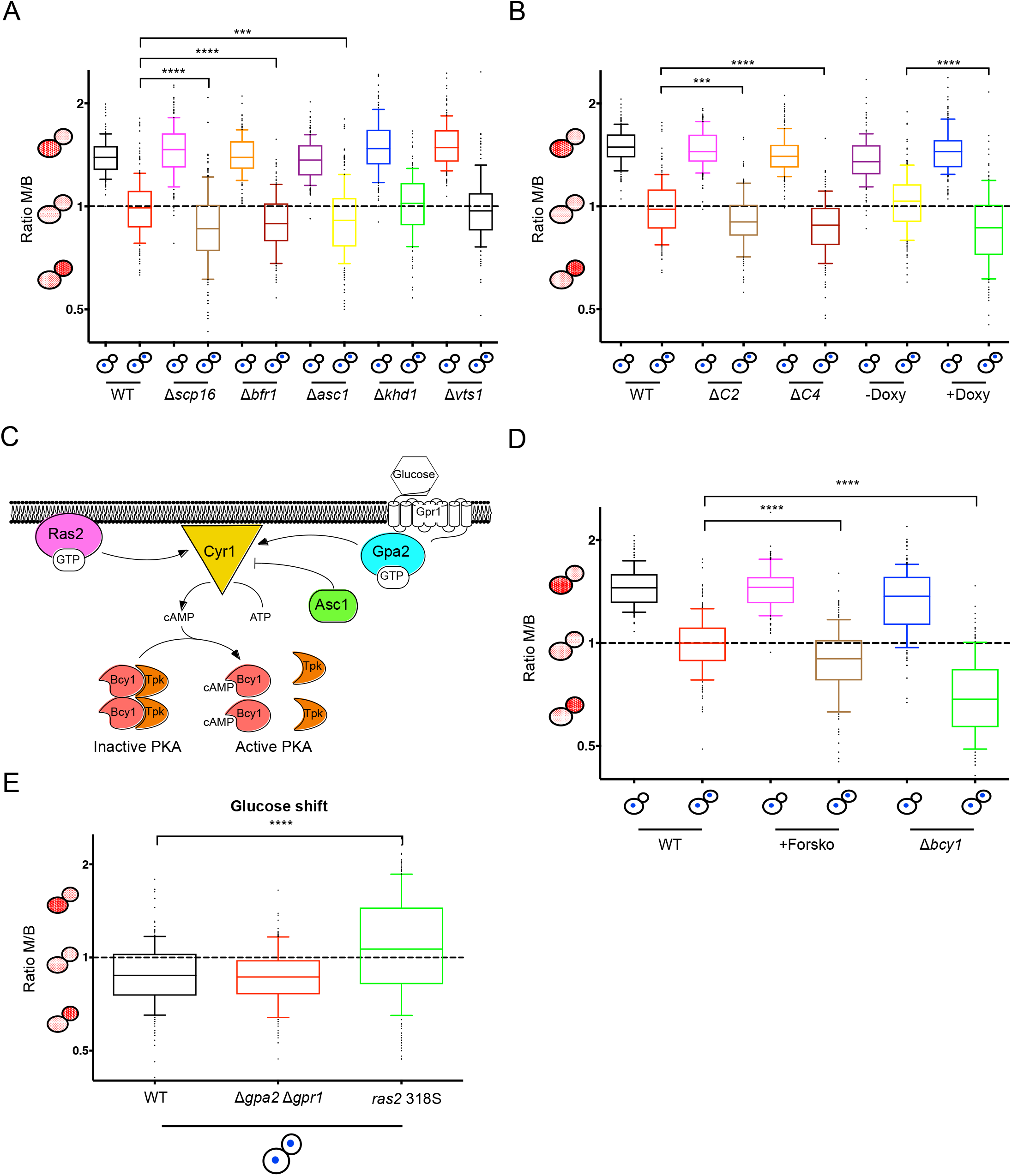
Loss of the Scp160/Bfr1/Asc1 complex and PKA cause enrichment of HXT2 mRNA in the bud after MAT. (**A**) Deletion of *SCP160, ASC1,* or *BFR1* increases HXT2 mRNA signal in the bud. FISH experiment with deletions in various RNA binding proteins. (**B**) The effect of *Δscp160* on HXT2 mRNA localization is independent of its increased ploidy. Scp160 truncations that lack either the last two (ΔC2) or the last four (ΔC4) KH domains or a Tet-off *SCP160* construct, in which expression is blocked by the addition of doxycycline (Doxy) confirm the phenotype observed in (**A**). (**C**) Schematic depiction of the glucose responsiveness pathway. (**D**) Hyperactivation of PKA drives accumulation of HXT2 mRNA in the bud. Treatment of cells with forskolin (+Forsko) or using a *Δbcy1* strain, causes HXT2 mRNA to be bud-enriched. (**E**) HXT2 mRNA enrichment upon refeeding is Ras2-dependent. Quantification of FISH experiments in (**A, B, D** and **E**) was performed as in Fig. 1C. ****: P < 0.0001; ***: P < 0.0005 in a two-tailed, unpaired t-Test.

The association of Scp160 with polysomes depends on Bfr1 and Asc1 (Baum et al., 2004, Lang, Li et al., 2001). Deletion of either *BFR1* or *ASC1* showed a phenotype indistinguishable from *scp160* mutants or depletion (Fig. 3A). The phenotype was specific because deletion of two other RNA binding proteins involved in mRNA localization, Khd1 and Vts1, did not show HXT2 mRNA accumulation in the bud (Fig. 3A).

Scp160 and Asc1 are part of the SESA complex, which functions in translational control of a subset of mRNAs (Sezen et al., 2009). Yet, deletion of another component of this complex, Eap1, had no effect on HXT2 mRNA localization, indicating that Scp160, Bfr1, and Asc1 act independently of the SESA complex in this process (Fig. S3B). Taken together, our results demonstrate that lack of Scp160, Bfr1 and Asc1 causes asymmetric HXT2 mRNA distribution after MAT.

The HXT2 mRNA enrichment in the bud in *scp160, bfr1* and *asc1* mutants was rather unexpected and very surprising. Therefore, we decided to further explore the mechanism and function of HXT2 mRNA enrichment in the bud after MAT.

### Increase of cAMP levels and PKA activity promote asymmetric distribution of HXT2 mRNA after metaphase/anaphase transition

Asc1 is the functional orthologue of mammalian RACK1 and was reported to function as a G-protein β-subunit coupled to glucose responsiveness (Zeller, Parnell et al., 2007) (Fig. 3C). Since Hxt2 is a glucose transporter, it is conceivable that Asc1’s function in repressing adenylate cyclase to keep cAMP levels low, would prevent HXT2 mRNA accumulation in the bud. Conversely, in this scenario, raising cAMP levels should drive HXT2 mRNA accumulation in the bud. To test this idea, we treated cells with forskolin, an activator of adenylate cyclase. As expected, forskolin did not affect HXT2 mRNA localization early in the cell cycle (Fig. 3D). However, we observed an increase in HXT2 mRNA signal in the bud comparable with that observed in the *Δasc1, Δbfr1* and *scp160* mutants. High cAMP levels cause the activation of PKA by binding to the inhibitory subunit Bcy1, which dissociates from PKA in the cAMP-bound form (Fig. 3C). We tested whether PKA signaling was involved in the HXT2 mRNA bud enrichment by deleting *BCY1.* In fact, HXT2 mRNA enrichment was even more pronounced in *Δbcy1* cells (Fig. 3D), indicating that indeed activation of PKA was responsible for HXT2 mRNA enrichment in the bud after mitosis.

Rather than using mutants to up-regulate cAMP/PKA signaling constitutively, we sought an approach to recapitulate transient cAMP production and hence transiently activate PKA. When starved cells are fed with glucose, they transiently increase cAMP levels (Jiang, Davis et al., 1998). We decided to use this regime to explore the changes in HXT2 mRNA localization upon transient PKA activation. We starved cells for 2 h and then shifted them to glucose-rich medium for 30 min prior to FISH analysis. HXT2 mRNA was enriched in the bud after MAT (Fig. 3E), demonstrating that indeed the cAMP/PKA pathway is responsible for the enrichment of HXT2 mRNA in the bud. This enrichment was also confirmed by single molecule FISH (smFISH) (Fig. S3C). The cAMP/PKA pathway can be activated through G-protein coupled receptor activation or Ras (Jiang et al., 1998, Xue, Batlle et al., 1998). To determine the upstream component of the adenylate cyclase, we deleted the receptor, *GPR1,* and the α-subunit *GPA2,* and used a strain carrying the *ras2^318S^* mutation (Jiang et al., 1998). While loss of the G-protein-coupled receptor branch did not interfere with HXT2 mRNA localization after refeeding, the *ras2^318S^* mutant was defective in HXT2 mRNA enrichment in the bud after MAT (Fig. 3E). We conclude that HXT2 mRNA enrichment in the bud after MAT is dependent on the Ras/cAMP/PKA signaling pathway. Since PKA is also involved in *HXT2* transcriptional activation (Kim & Johnston, 2006), we wondered whether transcription could contribute to the asymmetric HXT2 mRNA localization. When we blocked transcription with 1,10 phenanthroline after glucose shift, we did not observe an HXT2 mRNA enrichment in the bud after MAT (Fig. S2A and B), indicating that *de novo* synthesis is required for the asymmetric mRNA distribution under these conditions.

### Hexose transporter mRNAs are retained in the mother but not all of them are enriched in the bud when released from starvation

Seventeen *HXT* genes are encoded in the *S. cerevisiae* genome. We aimed to determine whether changes in mRNA localization are a common feature among HXTs. We concentrated on the 3 most important Hxts besides Hxt2: the low affinity transporters Hxt1 and Hxt3 and the high affinity transporter Hxt4 (Fig. 4A). Hxt2 can adopt at least two conformations that have been proposed to reflect high affinity and low affinity transporter states (Perez, Luyten et al., 2005, Reifenberger, Boles et al., 1997). Therefore, Hxt2 is special in that it cannot be clearly assigned to the low or high affinity transporter group. HXT1, HXT3 and HXT4 mRNAs were each retained in the mother early in the cell cycle and equally distributed after mitosis (Fig. 4B and C). However, HXT1, HXT3 and HXT4 mRNA were less responsive than HXT2 mRNA to high PKA levels in *Δbcy1* cells. HXT3 mRNA did not show any enrichment in the daughter cell (Fig. 4D). In a *Δbcy1* strain, PKA is constitutively hyperactive. To investigate HXT mRNA localization in a more physiologically relevant context, we again shifted cells from starvation to glucose to induce a short cAMP peak, which in turn is sufficient to activate PKA. Under those conditions, HXT1, HXT3 and HXT4 mRNA did not show any enrichment in the daughter cell after mitosis, while HXT2 mRNA was still enriched (Fig. 4E).

**Figure 4:**
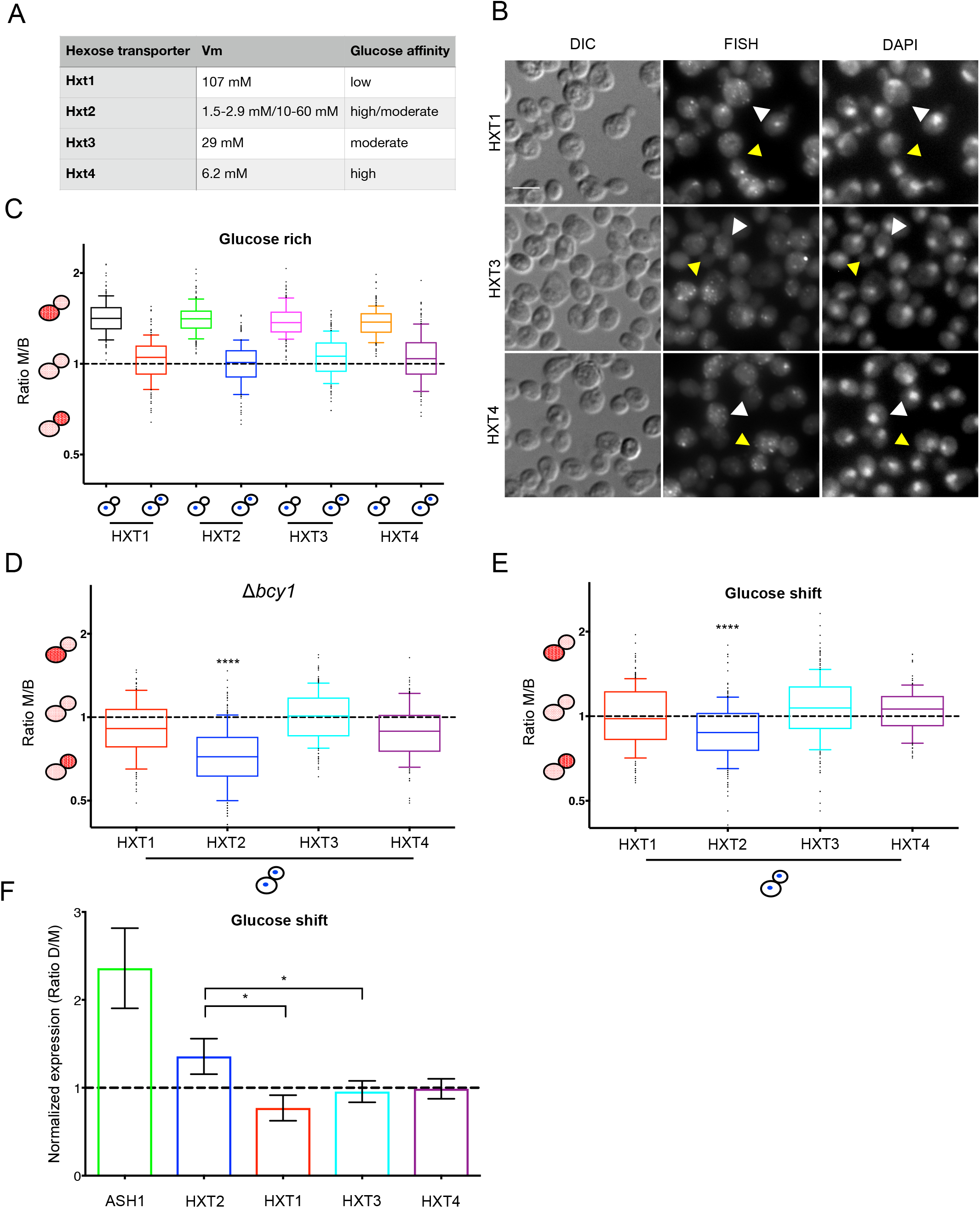
HXT mRNAs are retained in the mother before MAT, but only HXT2 mRNA is enriched in the bud after MAT upon refeeding. (**A**) Table of the glucose affinity of the four most important hexose transporters. (**B**) FISH of HXT1, 3 and 4 under glucose rich conditions. White arrowheads depict cells before, yellow arrowheads cells after MAT. (**C**) Quantification of (**B**). (**D**) Hyperactive PKA promotes enrichment of all HXTs after MAT. (**E**) HXT2 mRNA is the only HXT mRNA showing enrichment in the bud after MAT upon transient PKA activation. (**C-E**) Quantification of FISH experiments performed as described in Fig. 1C. **** P < 0.0001 in a two-tailed, unpaired t-Test. (**F**) Elutriation confirms enrichment of HXT2 mRNA in daughter cells upon refeeding. Daughters and mothers were separated through elutriation. qPCR was performed to determine the relative levels of different HXTs and ASH1 mRNA in both fractions. The mean of three independent experiments is plotted; bars represent standard deviation. *: P < 0.05 in a two-tailed, unpaired t-Test. Scale bar in (**B**); 5 μm.

To corroborate our findings, we separated young, newborn daughters from mother cells through elutriation during which we switched from no glucose to glucose-rich medium. We then extracted the total mRNA from the different cell populations and performed qPCR for HXTs. As a control, we used ASH1 mRNA, which is known to be enriched in daughter cells. ASH1 mRNA was about 2.3-fold enriched in daughter cells, a value which is in good agreement with the 2-2.2-fold increase reported previously (Takizawa, DeRisi et al., 2000). Among the HXT mRNAs, HXT2 was the only mRNA that accumulated in daughter cells (Fig. 4F). Taken together, retention in the mother prior to entering mitosis appears to be a general feature among the most important hexose transporter mRNAs, while the enrichment in the bud late in the cell cycle after release from starvation is specific for HXT2 mRNA.

### The spindle positioning protein Kar9 and nuclear segregation are essential for HXT2 mRNA bud enrichment under glucose shift conditions

Even though *de novo* HXT2 mRNA synthesis is required for bud enrichment (Fig. S2A and B), overexpression of HXT2 mRNA by itself was not sufficient to promote asymmetric mRNA distribution (Fig. S2C and D). Therefore, an additional layer of regulation must be required.

It is conceivable that the enrichment of HXT2 mRNA in the bud late in the cell-cycle could be dependent on three not mutually exclusive mechanisms: active transport of the mRNA into the bud, diffusion and retention, and selective degradation in the mother cell. P-bodies are the primary site of mRNA decay in yeast, and the exonuclease Xrn1 is responsible for the degradation of most mRNAs (Muhlrad, Decker et al., 1994). Therefore, we determined HXT2 mRNA localization after shift from starvation to glucose containing medium in *Δxrn1* cells. We could not detect a significant difference in mRNA localization when compared to wild-type cells (Fig. 5A). Thus, localized mRNA decay is an unlikely mechanism for the asymmetric HXT2 mRNA localization. To distinguish between active and passive transport, we repeated the treatment of cells with benomyl and LatA under the glucose shift conditions. Both benomyl and LatA treatment abrogated the HXT2 mRNA enrichment. (Fig. 5B). In addition, inhibition of the Arp2/3 complex by CK-666 inhibited HXT2 mRNA accumulation (Fig. 5B). Our data suggest that actin and MTs both contribute to the asymmetric distribution of HXT2 mRNA. Accordingly, we sought a cellular process involving both MTs and actin. Spindle positioning and spindle pole body (SPB) movement to the bud is such a process (Miller, Matheos et al., 1999). A key component in this process is the APC ortholog Kar9. Deleting *KAR9* reduced HXT2 mRNA asymmetric localization to the bud after MAT upon refeeding (Fig. 5C and D). In a *Δkar9* mutant, the old SPB is no longer preferentially segregated to the bud/daughter cell, and oftentimes both nuclei are retained in the mother cell after mitosis (Miller & Rose, 1998, Pereira, Tanaka et al., 2001). In large budded-cells after MAT, we observed equal HXT2 mRNA distribution when mother and daughter received a nucleus. In contrast, in cells in which both nuclei remained in the mother, HXT2 mRNA was likewise retained (Fig. 5C and D). Thus, Kar9 and potentially nuclear segregation into the bud play an essential role in asymmetric HXT2 mRNA distribution.

**Figure 5:**
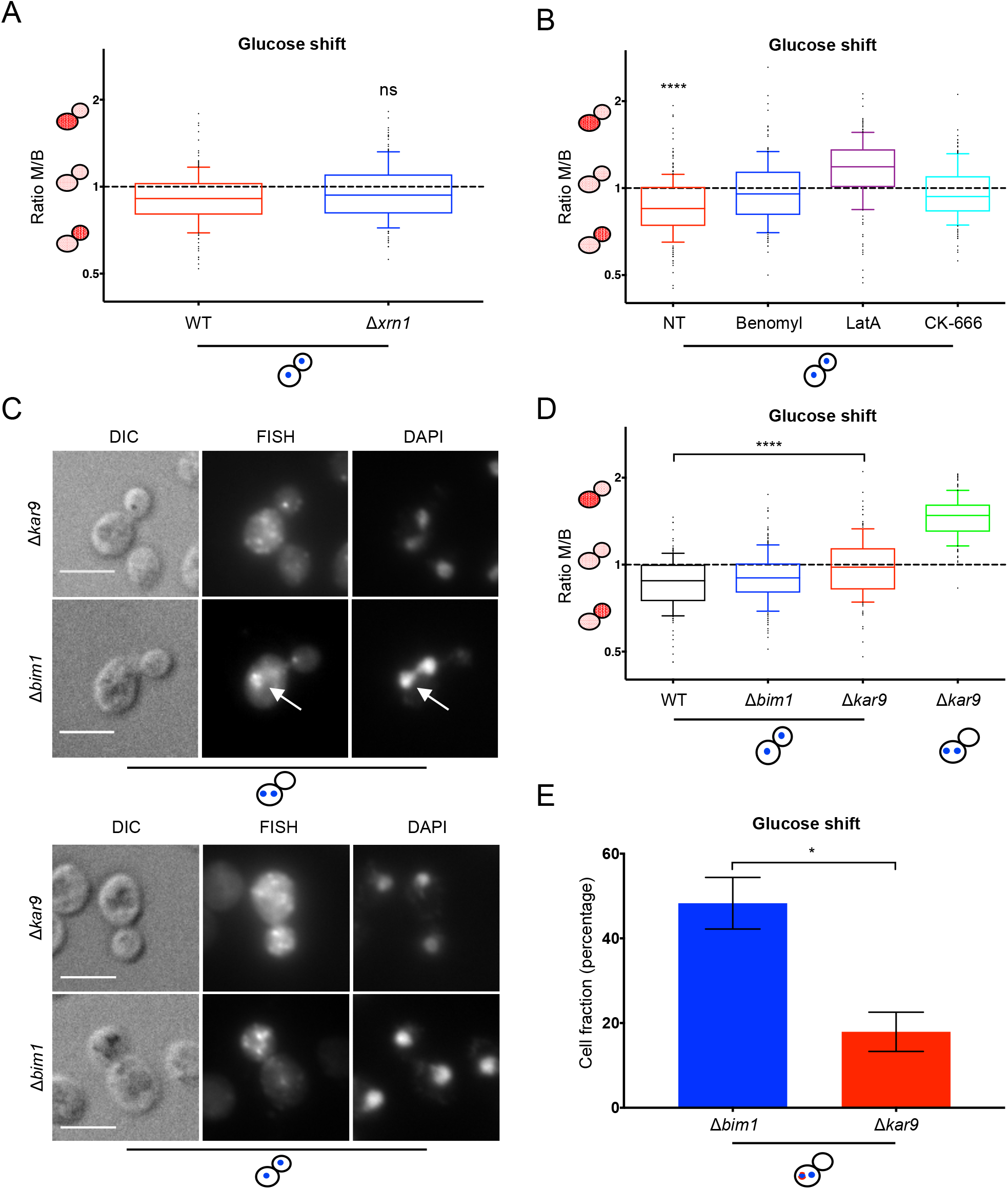
HXT2 mRNA enrichment in the bud depends on Kar9 and nuclear segregation. (**A**) RNA degradation is not essential for asymmetric HXT2 mRNA localization. FISH of wild-type and *Δxrn1* cells. (**B**) The cytoskeleton contributes to HXT2 mRNA enrichment in the bud. Cells were treated upon glucose addition with either 120 mg/ml benomyl for 15 min, 30 mg/ml latrunculin A for 30 min or 200 μM CK-666 for 30 min. HXT2 mRNA was detect by FISH. (**C**) Asymmetric loading of HXT2 mRNA onto the nuclear envelope and enrichment in the bud requires Kar9 but not Bim1 function. HXT2 mRNA FISH with *Δbim1* and *Δkar9* strains. Arrows point to a nucleus with HXT2 signal. (**D**) Quantification of the daughter cell enrichment from data displayed in (**C**). (**E**) Quantification of the HXT2 mRNA signal in cells with two nuclei in the mother cell from the experiment depicted in (**C**). Percentage of cells with asymmetric HXT2 mRNA staining is displayed. *: P < 0.05 in a two-tailed, unpaired t-Test. Quantification of the FISH experiments in (**A, B** and **D**) was performed as in Fig. 1C. ****: P < 0.0001 in a two-tailed, unpaired t-Test. Scale bars in (**C**) correspond to 5 μm.

### Kar9 is required for asymmetric distribution of HXT2 mRNA on the nuclear envelope

To corroborate our results, we deleted another component of the spindle positioning pathway, the EB1 homolog Bim1 (Lee, Tirnauer et al., 2000). In *Δbim1* cells, in which one of the nuclei reached the bud, HXT2 mRNA asymmetric distribution was largely unaffected (Fig. 5C and D). However, when both nuclei were retained in the mother, HXT2 mRNA was likewise retained. Our results are consistent with HXT2 mRNA requiring nuclear segregation into the bud for its asymmetric enrichment after glucose shift. Moreover, this process appears to be dependent on the spindle positioning pathway. Yet, the effect of *Δkar9* and *Δbim1* on HXT2 mRNA localization in mothers containing two nuclei was different. While in *Δbim1* cells HXT2 mRNA was preferentially localized to one of two nuclei, in *Δkar9* cells, the mRNA was not closely associated with a particular nucleus, and often not even recruited to the nuclear envelope (Fig. 5C and E). These results suggest that Kar9 is required for asymmetric loading of HXT2 mRNA on the nuclear envelope.

### The nuclear pore components Nup2 and Mlp1/2 are involved the asymmetric localization of HXT2 mRNA

We have shown above that PKA activation and nuclear segregation in the bud are prerequisites for the asymmetric distribution of HXT2 mRNA upon refeeding. Even though Kar9 is phosphorylated, this phosphorylation event has not been attributed to PKA action. We consequently hypothesized that PKA might phosphorylate a protein at the nuclear envelope or the endoplasmic reticulum, which in its phosphorylated form might contribute to asymmetric HXT2 mRNA localization. To test this hypothesis, we searched databases for ER/nuclear envelope localized PKA substrates (Table S1). Within this list, we found two components of the nuclear pore complex (NPC), which were promising candidates for asymmetric localization of mRNA - Nup2, which is involved in NPC inheritance (Suresh, Markossian et al., 2017) and the FG-repeat containing central core component Nup53. Indeed, when we deleted *NUP2,* the asymmetric localization of HXT2 mRNA to the bud was dampened (Fig. 6A and B). In contrast, a deletion mutant of *NUP53* and its paralog *NUP59,* did not affect the asymmetric HXT2 mRNA distribution (Fig. 6A and B). The *S. cerevisiae* NPC components Mlp1/2 were shown to be enriched at NPC insertion sites during anaphase at the old SPB that enters the bud (Ruthnick, Neuner et al., 2017). Deletion of *MLP1/2* resulted in a similar defect on HXT2 mRNA enrichment to *Δnup2* (Fig. 6A and B). Our data suggest a contribution by NPCs to the enrichment of HXT2 mRNA in anaphase in the bud upon refeeding, presumably through the NPC inheritance pathway. If our assumption was correct, we should be able to detect HXT2 mRNA on the nuclear envelope. To this end, we marked the nuclear envelope with Nup84-HA and performed FISH-IF. HXT2 mRNA was detected on the nuclear envelope in the bud (Fig. 6C).

**Figure 6:**
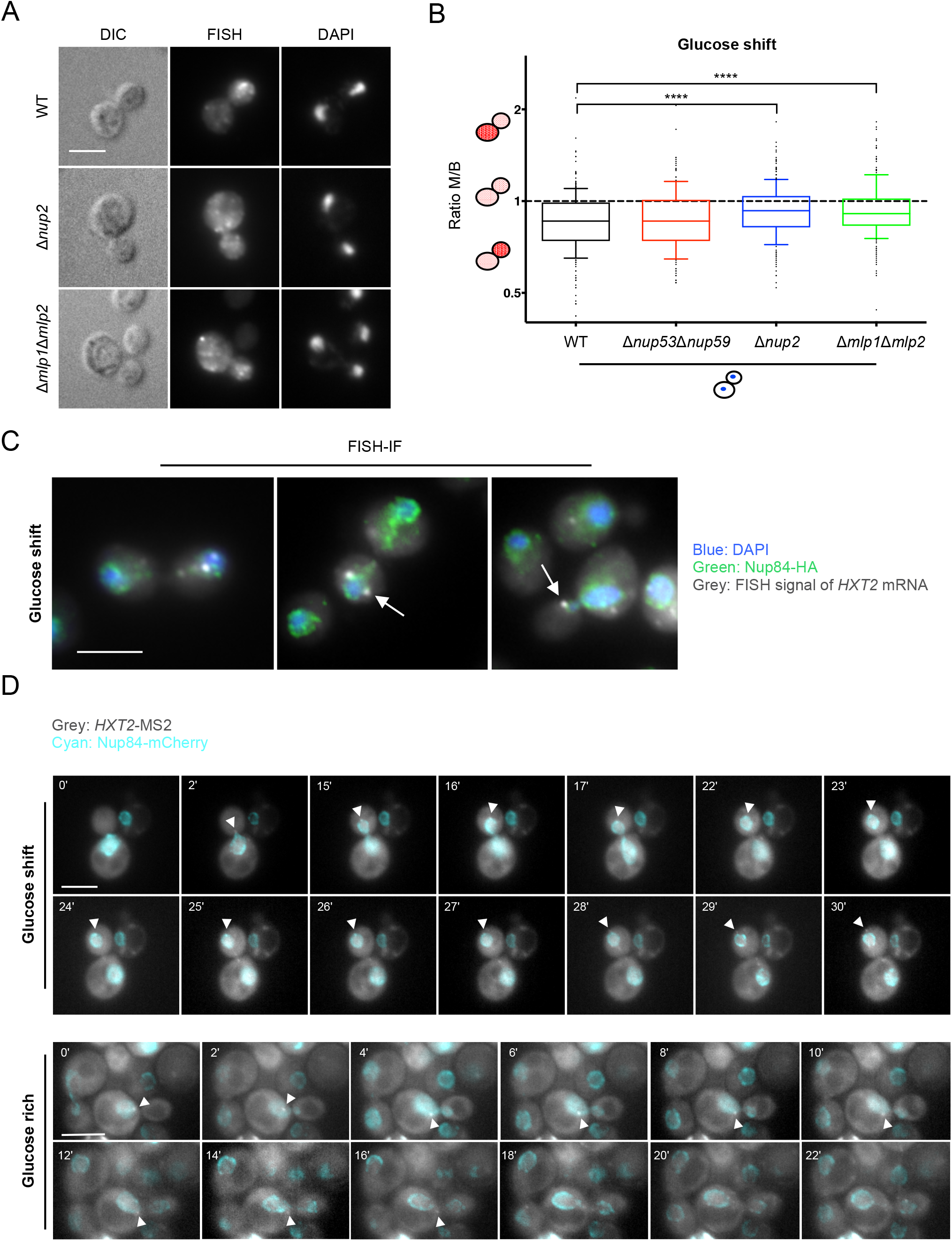
NPC components aid the enrichment of HXT2 mRNA in the bud upon refeeding. (**A**) Deletion of *NUP2* and *MLP1/2* reduced HXT2 mRNA enrichment in the bud. FISH under glucose shift conditions with strains in which different NPC components were deleted. (**B**) Quantification of (**A**). Quantification was performed as in Fig. 1C. ****: P < 0.0001 in a twotailed, unpaired t-Test. (**C**) HXT2 mRNA is on the nuclear envelope after MAT upon glucose shift. FISH-IF experiment with a probe against HXT2 mRNA and an antibody against the HA tag of the nuclear pore complex component Nup84-HA. White arrows point to HXT2 mRNA signals on the nuclear envelope. (**D**) HXT2 mRNA travels on the nuclear envelope into the bud during mitosis after glucose shift. Stills of time lapse movies. *HXT2* mRNA was visualized using the MS2-GFP system. Under glucose shift conditions *HXT2* mRNA is mostly localized to the bud together with the nucleus. Under glucose rich conditions *HXT2* mRNA moves mostly independent of the nucleus. Cyan: Nup84-mCherry; white arrowhead: *HXT2* mRNA-MS2 + MCP-GFP. Scale bars in (**A, C** and **D**) correspond to 5 μm.

To further corroborate our findings, we aimed to image movement of HXT2 mRNA during mitosis using an improved MS2-GFP based system (Tutucci, Vera et al., 2018). HXT2 mRNA was associated with the nucleus as it entered the bud in mitosis after refeeding (Fig. 6D, Movie S1). Such a movement was not observed in cells that did not experience glucose withdrawal (Fig. 6D, Movie S2). Taken together, our data provide strong evidence for an involvement of the nuclear envelope in the asymmetric HXT2 mRNA distribution during mitosis.

### HXT2 expression provides a growth advantage

Why would the yeast cell specifically enrich HXT2 mRNA into the bud following starvation? To gain insights into the biological function of this process, we used yeast strains in which all seventeen *HXT* genes had been deleted and which were kept alive by the expression of a single *HXT* gene under its endogenous promoter (Wieczorke, Krampe et al., 1999, Youk & van Oudenaarden, 2009). Those strains were either grown to stationary phase or cells were scraped off directly from a plate on which they were in a quiescent state. Serial dilutions were dropped on rich media plates. Growth of cells was assessed 2-3 days later. Cells expressing only *HXT2* grew better than cells expressing either only *HXT1, HXT3* or *HXT4* on a wide range of glucose concentrations (Fig. 7A and B, Fig. S4). Next, we tested whether HXT2 only strains would also do better when they were directly competing with the other HXT only strains in culture. To this end, we mixed stationary cultures of the 4 HXT only strains, collected a sample for the starting point and let them grow over night. Then we collected another sample and performed qPCR on the DNA. Under direct competition for resources, the HXT2 only strain grew faster and outcompeted the other strains (Fig. 7C and D). Collectively, our data suggest that asymmetric HXT2 mRNA distribution provides a growth advantage.

**Figure 7:**
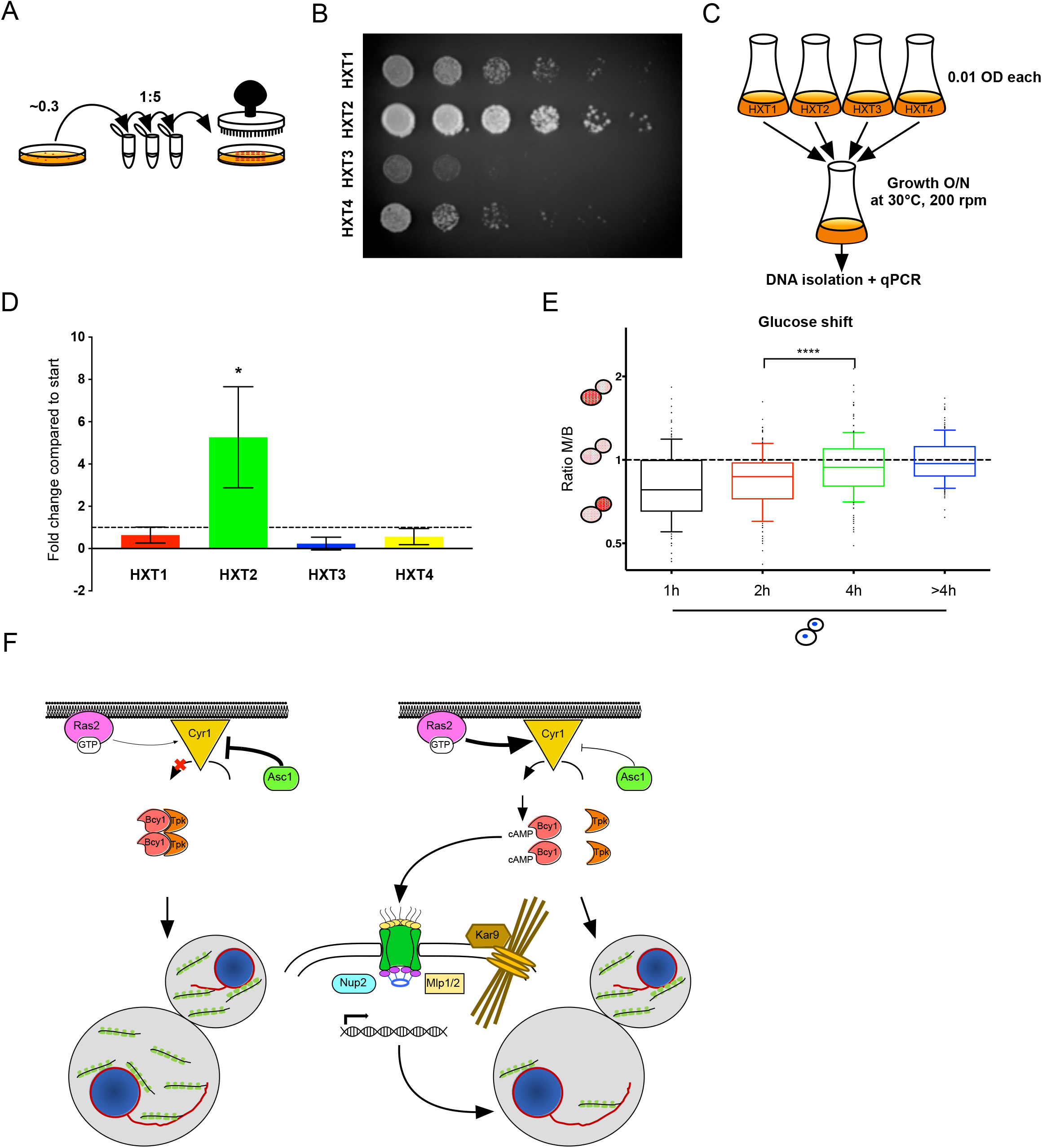
HXT2 mRNA enrichment in the bud under glucose shift conditions provides a growth advantage to daughter cells. (**A**) Schematic depiction of the growth assays. Cells expressing only one hexose transporter at a time were picked from a plate, serially diluted and plated on fresh YPD plates. (**B**) HXT2 only strains grow faster than the other HXT only strain. Growth was assessed 2-3 days after incubation at 30°C. (**C**) Workflow for the competition assay presented in (**D**). O/N cultures of HXT-only strains were diluted to the same OD_600_ and pooled. A sample was taken (‘start’), the rest of the culture incubated O/N at 30°C. Cells were harvested (‘end’), DNA was isolated from both samples, and the amount of *HXT* DNA was determined by qPCR. (**D**) Quantification of competition assay. qPCR with primers against either *HXT1, 2, 3* or *4* was performed. *HXT* gene abundance was normalized against actin. The results of three independent experiments are shown. Error bars represent standard deviation. Comparison of HXT-levels at the beginning and the end of growth shows that cells expressing only HXT2 outgrow other HXT-only cells in the range of 2 to 5-fold. *: P < 0.05 in a two-tailed, unpaired t-Test. (**E**) Daughter cells do not retain a memory of the previous stress situation encountered by their mothers. Cells were re-fed after starvation and analyzed for enrichment of HXT2 mRNA via FISH at different time points. Quantification was performed as in Fig. 1C. ****: P < 0.0001 in a two-tailed, unpaired t-Test. (F) Proposed model for the regulation of the asymmetric HXT2 mRNA distribution when cells come out of starvation or leave the quiescent state. For details, please see text.

### The cell does not retain a long-lasting memory for HXT2 mRNA localization

We envisage that the increase in HXT2 mRNA in newborn daughters, will allow those daughters to take up more glucose and therefore grow faster than their counterparts that do not increase HXT2 expression. In this scenario, the daughters with high HXT2 will also reach the critical size earlier to enter the cell cycle and become a mother. We wondered, whether asymmetric HXT2 mRNA distribution would be retained in future generations in order not lose the competitive edge. In other words, we asked whether high HXT2 daughters remember that they received more HXT2 and will therefore also asymmetrically distribute the HXT2 mRNA during the next cell division, even though the daughters have not experienced nutrient limiting conditions themselves? Alternatively, there is no memory of the previous events transmitted to the daughters because entering the cell cycle faster comes at an expense in that errors that occurred during previous cell cycle may not be corrected. To distinguish between the two possibilities, we performed a glucose shift experiment and took samples at different time-points after the shift. FISH revealed that a significant enrichment in the daughter was observed only up to 2 h after shift (Fig. 7E). Considering the lag phase upon resuming the cell division cycle, these results indicate that a newborn daughter does not retain a memory in terms of HXT2 mRNA localization of its mother’s starvation experience, when it becomes a mother itself.

## Discussion

We have identified a mechanism in *S. cerevisiae* by which the mRNA of the glucose transporter Hxt2 is specifically enriched in the daughter cell when cells are coming out of quiescence or starvation. We assume that this enrichment provides a growth advantage to these young daughters as they are potentially able to take up more glucose and thus increase in size faster than daughters with less Hxt2. This mechanism allows those daughters to outcompete cells that cannot mount a similar response and provides an ecological advantage. This partitioning is not shared by other major glucose transporters, underscoring a unique role of Hxt2. Hxt2 differs from the other glucose transporters in that it can act as both high affinity and low affinity transporter (Perez et al., 2005), rendering it particularly suitable for such a regulation. We demonstrate that accumulation of HXT2 mRNA is dependent on the Ras/cAMP/PKA pathway and the nuclear envelope. We speculate that the short cAMP peak upon refeeding provides sufficient PKA activity to phosphorylate targets on the nuclear envelope such as Nup2 (Figure 7F). Our data are consistent with a model in which PKA phosphorylates the NPC component Nup2. This would then in turn allow the piggy-back of HXT2 mRNA into the daughter cell. Whether this piggyback is through the NPC inheritance pathway, direct interaction with Nup2 or correlates to NPC function remains to be determined. However, we favor at this point the involvement of the NPC inheritance pathway rather than NPC function or direct binding to Nup2 based on the observation that deletion of another NPC PKA target, the FG repeat protein Nup53 and its paralog Nup59, did not affect HXT2 mRNA localization. These data argue against NPC function per se being essential for the asymmetric mRNA localization. In contrast, Mlp1/2, which are NPC components that are enriched in the daughter cell during mitosis are likewise contributing to the accumulation of HXT2mRNA in the daughter cell (Ruthnick et al., 2017). Importantly, NPCs have been shown to accumulate around the yeast centrosome (SPB) presumably through a mechanism involving Mlp2 and Nup2 (Niepel, Strambio-de-Castillia et al., 2005, Suresh et al., 2017, Winey, Yarar et al., 1997).

Our data provide strong evidence that movement of the nucleus into the bud is critical for asymmetric enrichment of HXT mRNA. Both actin and microtubules as well as the yeast EB1 and APC homologs, Bim1 and Kar9, respectively, are essential for this process.

Yet, it appears that Kar9 has an additional role in that it may help to generate the asymmetry on the nuclear envelope. While in a *Δbim1* background HXT2 mRNA was present preferentially only on one of two nuclei that were retained in the mother, in *Δkar9* this asymmetry was abolished. During spindle formation early in the cell-cycle Kar9 becomes enriched on the old SPB, which will move into the bud during mitosis in a process dependent on both actin and microtubules (Cepeda-Garcia, Delgehyr et al., 2010, Liakopoulos, Kusch et al., 2003, Maekawa & Schiebel, 2004). During this process, Kar9 translocates from the SPB to the MT (+) ends in the bud by binding to Bim1 (Huls, Storchova et al., 2012). Therefore, it is likely that Kar9 asymmetry is instructive for HXT2 mRNA localization. However, the underlying mechanism that controls Kar9 asymmetry remains unclear and highly controversial (Juanes, Twyman et al., 2013).

We also found that *HXT2* transcription contributed to the asymmetric enrichment in the bud upon refeeding. However, transcription per se was not sufficient because *HXT2* overexpression did not influence mRNA localization. The *HXT2* transcription during mitosis in the glucose shift experiments might be driven by the same Ras and cAMP-dependent peak of PKA activity (Kim & Johnston, 2006) that drives asymmetric mRNA localization. This scenario would elegantly couple gene transcription to asymmetric distribution of its mRNA.

We generally observed that HXT mRNA was retained in the mother cell before MAT. We assume that the mRNAs are engaged with ribosomes and membrane bound. This retention was also coupled, at least for HXT2, with translational stalling. The mechanism by which this is achieved remains unclear but does not appear to involve SRP. We suppose that at least a part of the stalling of the ribosomes is due to some cis-elements on the mRNA itself. However, since this is a regulated process, trans-acting factors are likely to play a role as well. How these stalled ribosomes would escape ribosome quality control needs also be investigated. Further studies are needed to elucidate the retention mechanism of HXT2 retention.

We propose a model in which the asymmetric HXT2 mRNA distribution upon refeeding allows the daughters to uptake more glucose and therefore grow faster than daughters with less HXT2 mRNA. Since cell size is one of the hallmarks for entering a new cell cycle, the faster growing daughters will start a new division cycle before the others. The faster entry into the cell cycle appears to be sufficient to out compete the slower growing cells. Yet, the faster growing daughters do not retain an obvious memory of the glucose limitation and refeeding because they do not provide more HXT2 mRNA for their daughters. This finding is consistent with the observation that a temporal cAMP/PKA activity peak is sufficient to drive the mRNA enrichment, and hence a memory may not be needed. Moreover, it may not be advantageous for the cells to always enter the cell division cycle as fast as possible. Before passing START, the cells usually check whether the last cell division was successful on multiple levels and aim to repair defects that may have occurred.

Therefore, the time between two cell division cycles needs to be balanced between speed and fidelity.

Maintaining such a balance is in contrast to what likely happens in cancer cells. In numerous cancer types, glucose transporters (GLUTs), in particular GLUT1 and GLUT3, are upregulated and often linked to oncogenic RAS (Adekola, Rosen et al., 2012). Like HXTs in yeast, GLUTs are localized in polarized fashion in epithelial cells. Moreover, GLUT3 expression is up-regulated by cAMP in a breast cancer cell line (Meneses, Medina et al., 2008) and increased GLUT1 level have been correlated with poor prognosis in colorectal carcinomas (Sakashita, Aoyama et al., 2001). Thus, cancer cells may employ a similar strategy in terms of accelerating growth, but lack check points to monitor damage that occurred in the previous cell cycle. This may be different in cancer stem cells (CSCs), which are metabolically more similar to stem cells in that they can use glycolysis over OXPHOS and are often in hypoxic environments (Wong, Che et al., 2017). At least in embryonic stem cells GLUT1 and GLUT3 expression levels are correlated to maintaining pluripotency (Wu, Song et al., 2017, Zhang, Skamagki et al., 2017). Moreover, GLUT1 is asymmetrically localized in dividing lymphocytes and becomes enriched in the differentiating daughter (Chen, Kratchmarov et al., 2018). Therefore, the controlled expression of specific glucose transporter at the plasma membrane, is an important and conserved cellular feature to maintain competitiveness in difficult environments.

### Experimental Procedures

#### Yeast strains and growth conditions

Standard genetic techniques were employed throughout (Sherman, 1991). Unless otherwise noted, all genetic modifications were carried out chromosomally. Chromosomal tagging and deletions were performed as described (Gueldener, Heinisch et al., 2002, Janke, Magiera et al., 2004, Knop, Siegers et al., 1999). For amplification of C-terminal tagging by chromosomal integration pYM (Knop et al., 1999) and for deletions pUG plasmids (Gueldener et al., 2002) were used. For N-terminal tagging or promoter exchange by chromosomal integration pYM-N plasmids were used (Janke et al., 2004). Primers, strains and plasmids used in this study are listed in Supplementary Tables S2-4. The plasmids pMS449 and pMS450 carrying the ΔC2 and ΔC4 truncation of Scp160 were a kind gift from M. Seedorf (ZMBH Heidelberg, Germany). The plasmid pRJ1463 bearing Scp160 under the control of a Tet-off promoter was a kind gift from Ralf-Peter Jansen (Jansen, Niessing et al., 2014). For glucose rich conditions, yeast cells were grown in either YPD (1% w/v Bacto yeast extract, 2% w/v Bacto-peptone, 2% w/v dextrose) or selective medium (prepared as described (Kaiser, 1994)) at 30°C, 200 rpm. For glucose shift conditions, cultures were first grown over night in YPD, then washed two times in YP media without glucose diluted to an OD_600_ of 0.3 and further grown for 2 h in YP media without glucose. After this incubation, glucose was added to a final concentration of 2%. Generally, yeast cells were harvested at mid-log phase with an OD_600_ of 0.4-0.8. OD_600_ as determined with a UltroSpec 3100 pro Spectrophotometer (GE healthcare).

#### Life cell imaging

Yeast cells were grown in YPD or HC-Leu (2% dextrose, 1x adenine) to early log phase and either analyzed directly or first starved without dextrose for 2 h then supplemented with 2 % dextrose. The cells were taken up in HC-complete or HC-Leu medium and immobilized on 1% agar pads. Fluorescence was monitored with an Orca Flash 4.0 camera (Hamamatsu) mounted on an Axio M2 imager (Carl Zeiss) using a Plan Apochromat 63x/NA1.40 objective and filters for mCherry and GFP. ZEN software version 2.3 (blue edition) was used to acquire images (Carl Zeiss). Further Image processing was performed using Fiji (Schindelin, Arganda-Carreras et al., 2012). All pictures from the same experiment were treated equally.

#### Fluorescence in situ hybridization (FISH) and FISH combined with immunofluorescence (FISH-IF)

FISH and FISH-IF were performed as described previously (Kilchert & Spang, 2011, Takizawa, Sil et al., 1997). For IF, we used as primary antibodies anti-GFP (Torrey Pines, rabbit, polyclonal, 1:400), anti-HA (Covance, mouse, monoclonal, 1:400) and as secondary antibodies Alexa Fluor 488 goat-anti-rabbit or anti-mouse, respectively (Invitrogen, polyclonal, 1:400). Images were acquired with an Axiocam MRm camera mounted on an Axioplan2 fluorescence microscope using a Plan Apochromat 63x/NA1.40 objective and filters for eqFP611, DAPI and GFP. AxioVision software 3.1 to 4.8 was used to process images (Carl Zeiss).

##### FISH quantification - Image Processing and Analysis

Images were acquired on a Zeiss Axioplan2 microscope as described above. Segmentation and analysis were performed using a custom macro for Fiji (available upon request) carrying out the following steps: Based on the red (FISH) channel, the CLAHE (Zuiderveld, 1994) filter was executed to increase the local contrast with the goal of revealing enough contrast for being able to distinguish all cell bodies from the background. Parameters for the CLAHE filter were used with their defaults as provided by the plugin with a block size of 127, a default slope of 2 and 256 histogram bins. After filtering, the local thresholding method as described by (Phansalkar, 2011) was used to create a binary image marking the cell bodies (thresholding radius: 50, object size minimum: 50). To reduce artifacts in the segmentation the binary image was then processed with Fiji’s binary “Fill Holes” operation, followed by a “Close” operation. The resulting binary mask was then split into individual masks per cell by using the classical “Watershed” separation. As a last step for the segmentation individual ROIs (regions of interest) were created from this binary mask. After creating the ROIs, they were used to quantify intensities in the red (FISH) channel by measuring the mean gray value per round cell shape. With those results, fluorescence intensities for mothers and their buds were determined separately and finally the ratio of fluorescence intensity in the mother over bud was calculated. At least 50 mother cells and their corresponding buds from at least three biologically independent experiments were counted per condition.

#### Elutriation

Elutriation was performed in a Beckman elutriation system (Avanti J-26 XP centrifuge combined with a JE-5.0 elutriator rotor and a 200 ml elutriation chamber, as well as standard elutriation accessories) according to (Marbouty, Ermont et al., 2014) with modifications. For glucose shift experiments, first a colony was inoculated in 500 ml YPD medium and incubated overnight at 30°C. The next day, the cells were washed 2x with YP medium without glucose, then diluted to an OD_600_ ~1 in 2 x 1 l fresh YP medium without dextrose and further grown for 2 h. Dextrose was added to a final concentration of 2 % and the cultures were incubated for another 30 min at 30°C. The elutriation chamber was filled with the cell culture at a flow rate of ~25ml/min and a rotor speed of 3,000 rpm at 4°C. The flow rate was gradually increased until the cells reached the top of the chamber. Equilibrium was allowed to settle in the chamber for 1 h while switching medium to PBS. The flow through was checked for escaping cells by OD_600_ measurement. The flow rate was carefully increased by increments of 2 ml/min until the OD_600_ raised above 0.05, at which point cells were collected. The budding index of the recovered cells was checked by microscopy and only daughter cell fractions with less than 5 % budded cells were used. After collection of the daughter cells, the centrifuge was stopped and the remaining cells were collected, providing the mother cell fraction. The fractions were spun down. The pellets were frozen in liquid nitrogen and stored at −80°C.

#### Total RNA isolation

Total RNA was isolated from yeast essentially as described (Schmitt, Brown et al., 1990). Briefly, 10 OD_600_ of yeast cells were harvested after different treatments. Cells were resuspended in 1 ml AE buffer (50 mM NaOAc pH 5.2, 10 mM EDTA), 100 μl 20 % SDS and 1 ml PCI (125:24:1, pH 4.4) were added and the tube vortexed for 10 s. After incubation for 10 min at 65°C, tubes were chilled in liquid nitrogen for 30 s, thawed and then centrifuged (2 min, 16,000 x g, RT). The upper aqueous phase was transferred to a fresh tube and 1 ml PCI was added and vortexed 10 s. After phase separation by centrifugation, the upper aqueous phase was transferred to a fresh tube and 1/10 vol. of 3 M NaCl (pH 5.2) was added. The RNA was precipitated with 1 vol. 100 % EtOH at −80°C for at least 1 h. The precipitate was collected (30 min, 16,000 x g, 4°C), washed with 70 % EtOH and again centrifuged. The pellet was resuspended in 200 μl RNase-free water.

#### Quantitative RT-PCR

0.5-1 μg of total RNA was reversely transcribed with the Transcriptor reverse transcriptase kit (Roche, Cat# 03531287001), oligo-dTs and random hexamers. The mRNA levels were analyzed by SYBR green incorporation using the ABI StepOne Plus real-time PCR system (Applied Biosystems). Primers used in qRT-PCR are listed in Table S3.

#### Drop assays

Colonies were either directly taken from plate or grown overnight in liquid YP medium supplemented with 2% dextrose to stationary phase (OD_600_ ≥ 9). After adjusting to equal cell concentrations (OD_600_ ~ 0.3), five serial dilutions (1:5) were dropped onto YP plates supplemented with either 0.1, 0.5, 2, or 4 % glucose using a “frogger” stamp (custom-built). Plates were incubated for 2-3 days at 30°C and photographed for documentation.

#### Actin staining

Actin cytoskeleton staining was performed essentially as described previously (Adams & Pringle, 1991). Cells were grown over night and fixed with 4% formaldehyde for 30 min at RT under gentle agitation. Cells were washed twice with PBS containing 1 mg/ml BSA (3 min, 1,000 g, RT). The cell pellet was resuspended in 25 μl PBS containing 1 mg/ml BSA and 5 μl of rhodamine-phalloidin (Molecular Probes, 300 U/1.5 ml MeOH). After incubation for 1 h at RT in the dark, cells were washed three times and resuspended in 500 μl PBS containing 1 mg/ml BSA. An aliquot was allowed to settle for 30 min on polyethyleneimine-treated multiwell slides. The slides were washed briefly in PBS, Citifluor AF1 was added and the coverslips were sealed with nail-polish. Slides were stored at −20°C. Rhodamine fluorescence was observed by epifluorescence microscopy using the Cy3 channel on an Axioplan 2 fluorescence microscope from Zeiss. Pictures were taken with an Axiocam MRm camera using AxioVision software. Image processing was performed with Fiji.

#### Western Blotting

Nine ml of a mid-log grown culture were taken, immediately treated with cold trichloroacetic acid (10% final concentration) and incubated on ice for at least 5 min. Yeast extracts were prepared as described (Stracka, Jozefczuk et al., 2014). The protein concentration was determined using the DC Protein Assay (Bio-Rad) and the total lysate was analyzed by SDS-PAGE. The first antibody was rabbit anti-Scp160 (Weidner et al 2014, 1:1,000), secondary antibody was goat-anti-rabbit, HRP-conjugated (Pierce, polyclonal, 1:15,000). Enhanced Chemiluminescence solution (ECL; GE Healthcare) was used for detection.

#### PKA target prediction

To identify potential PKA targets, we intersected subcellular localization annotations and known phosphopeptides from yeast. The localization set of interest includes all UniProt entries for *S. cerevisiae* with at least one localization annotation of “endoplasmic reticulum”, “nucleus outer membrane”, or “nucleus envelope” (The UniProt, 2017). Yeast phosphopeptides were identified from the ISB Library of the PeptideAtlas Project (Desiere, Deutsch et al., 2006) “The PeptideAtlas Project”, Nucleic Acids Research 34, D655-D658]. Custom Python scripts were used to identify exact matches between phosphopeptides of the yeast PeptideAtlas and the protein sequences of the UniProt entries filtered by localization.

## Acknowledgements

We thank J. Broach, E. Boles, S. Rospert, R.P. Jansen, M. Seedorf, R. Singer, and E. Schiebel for strains and reagents. We are grateful to E. Schiebel for discussions, and to I.G. Macara and K. Weis for critical comments on the manuscript. This work was supported by the Human Frontiers Science Program (RG0031/2009), the Swiss National Science Foundation (310030B-163480) and the University of Basel to AS and an EMBO longterm-fellowship (ALTF 289-2010) to SH.

The authors declare no competing financial interests.

---

**Figure S1:**
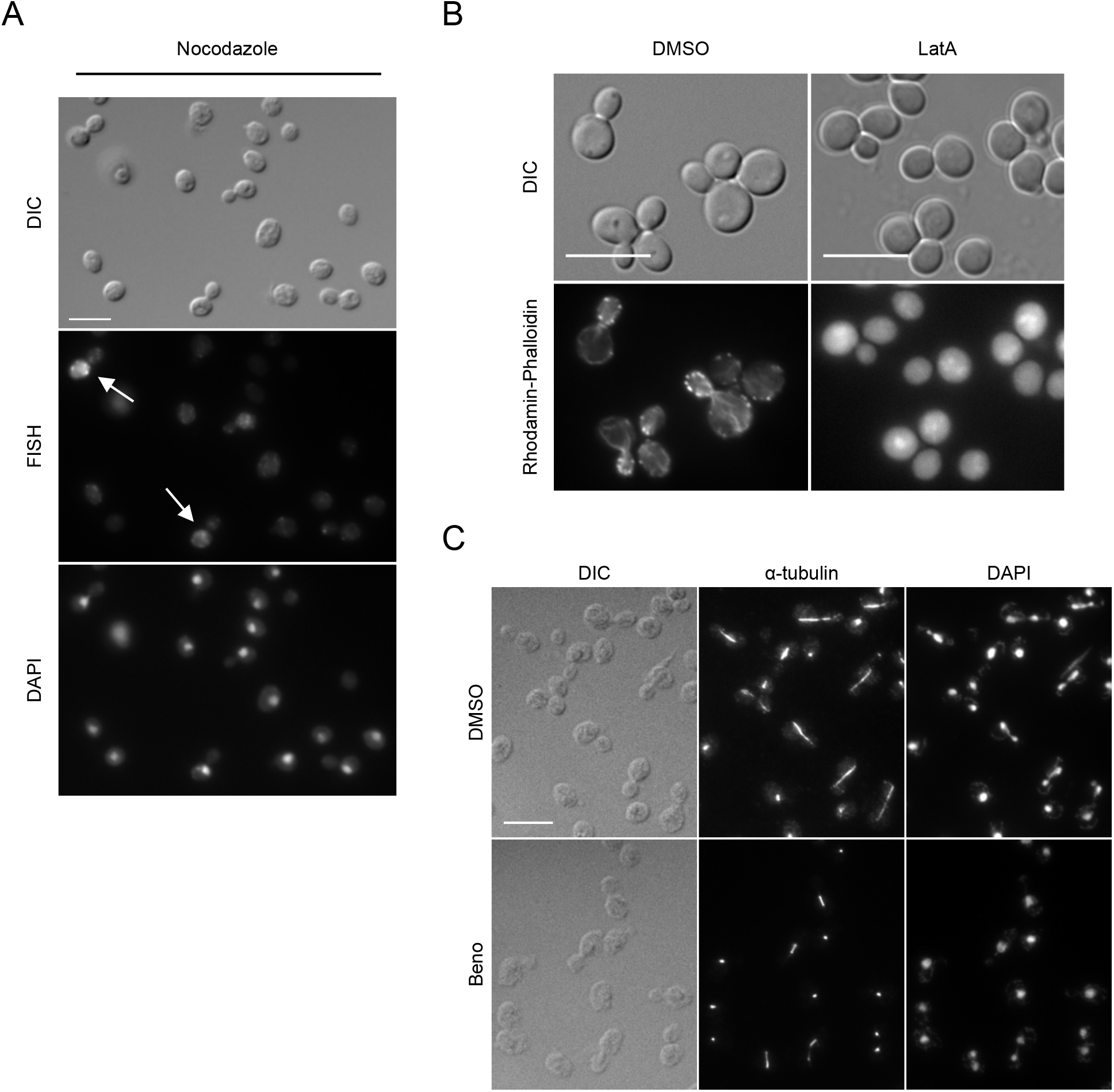
(**A**) *HXT2* mRNA release from the mother cell is coupled to cell cycle progression and nuclear segregation respectively. Cells arrested in G2/M phase with nocodazole show still retention of *HXT2* mRNA in the mother even in large budded cells (arrows). Cells were treated with 15 μg/ml for 3h, subsequently fixed and mRNA was visualized by FISH. (**B**) Rhodamin-phalloidin staining. Cells were either treated with 30 μg/ml Latrunculin A (LatA) or as a solvent control with DMSO for 30 min. After fixation, actin was stained with Rhodamin-phalloidin. LatA treated cells show no actin cables or patches anymore. (**C**) Benomyl treatment leads to the depolymerization of cytoplasmic microtubules but cells are still able to segregate the nuclei. Immunofluorescence of microtubules with an antibody against α-tubulin.

**Figure S2:**
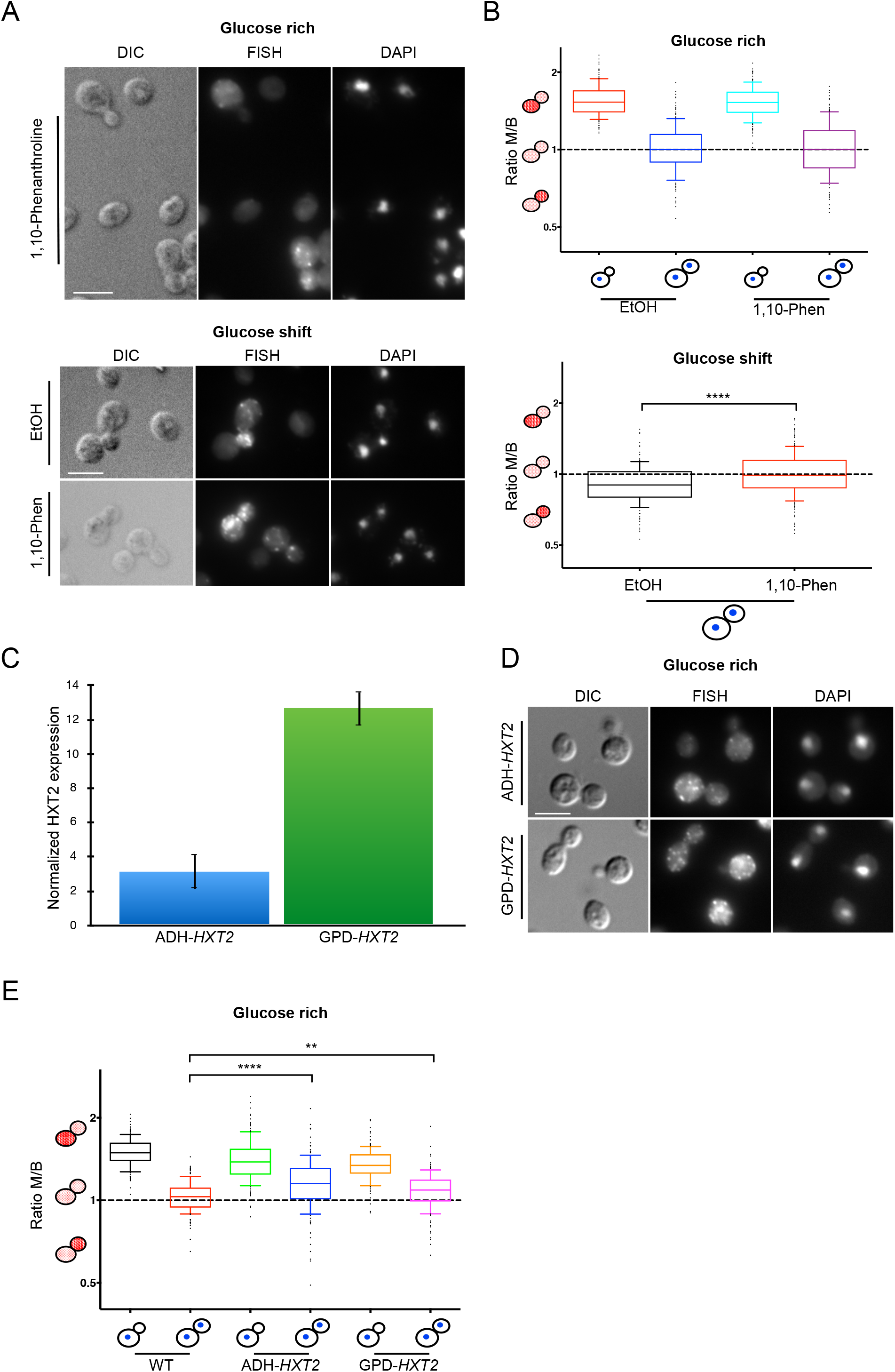
Inhibition of transcription with 1,10-phenanthroline (1,10-Phen) hampers the enrichment of *HXT2* mRNA in the bud under glucose shift conditions but not the equal distribution under glucose rich conditions. (**A**) Cells were treated with 100 μM 1,10-phenanthroline or the solvent control ethanol (EtOH) for 1h. (**B**) Quantification of (**A**). (C-E) Increased transcription of *HXT2* per se does not lead to its enrichment in the bud. ****: P < 0.0001 in a two-tailed, unpaired t-Test (**C**) Exchanging the 5’-UTR with either the ADH- or GPD-promotor leads to the overexpression of *HXT2.* Quantification of *HXT2* expression compared to WT with qPCR. (**D**) FISH experiments revealed that *HXT2* mRNA is still retained in the mother cell after MAT under glucose rich conditions when the native 5’-UTR is swapped for the ADH- or GPD-promotor. (**E**) Quantification of (**D**). ****: P < 0.0001 in a twotailed, unpaired t-Test. **: P < 0.05 in a two-tailed, unpaired t-Test.

**Figure S3:**
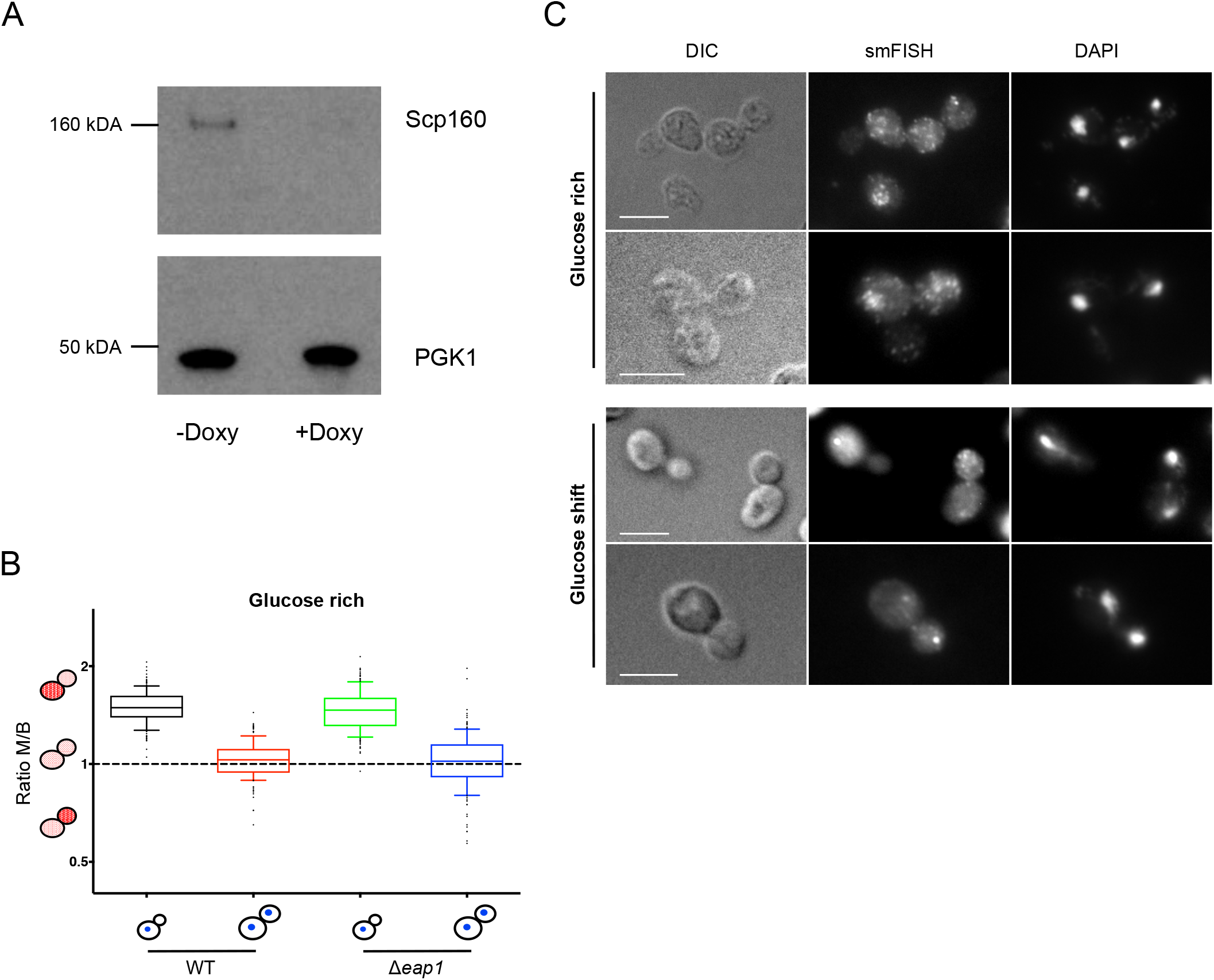
(**A**) Protein levels of Scp160 under the control of a Tet-off promotor markedly decrease when treated with 2 μg/ml Doxycycline (+Doxy) for 6h. PGK1 serves as loading control. (**B**) Deleting another component of the SESA complex does not have an influence on the localization of *HXT2* mRNA as compared to Scp160 or Asc1. (**C**) Single molecule FISH (smFISH) shows the same distribution pattern for *HXT2* mRNA under both, glucose rich and glucose shift conditions.

**Figure S4:**
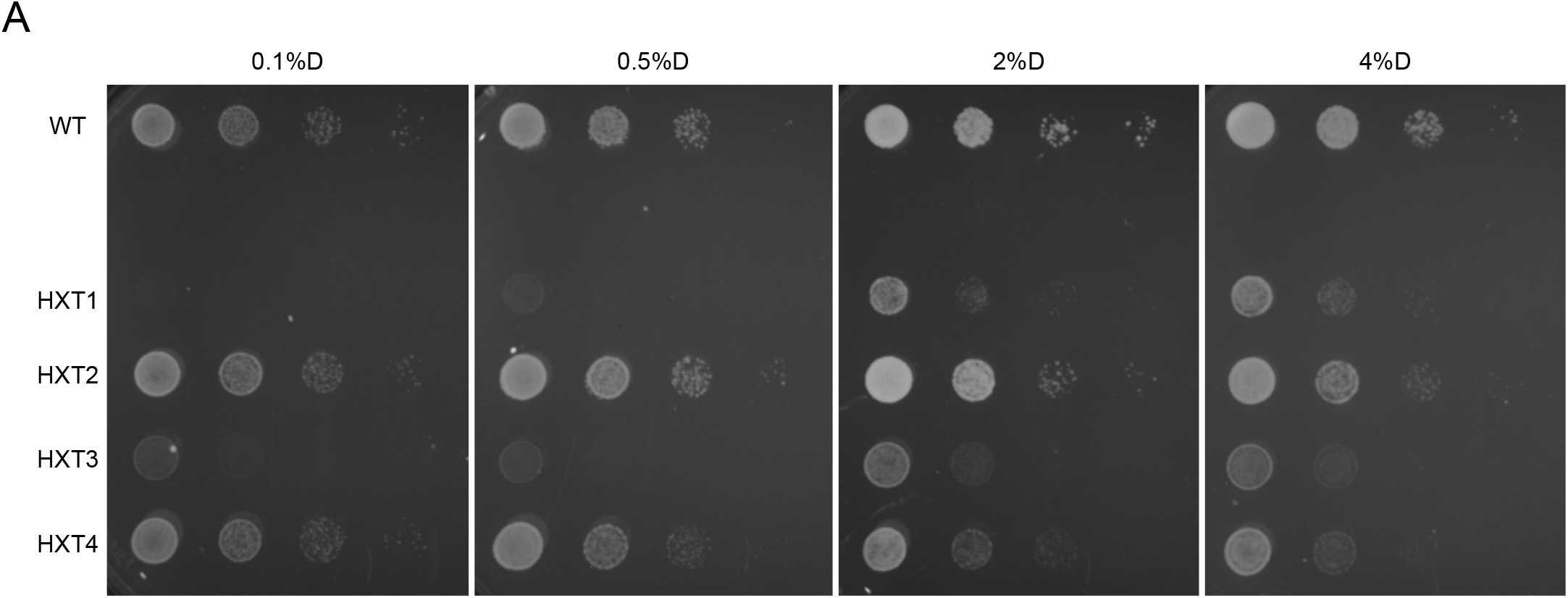
At different glucose concentrations HXT2 only cells still grow faster compared to other HXT only cells. Growth assay as described in 7A on plates with indicated glucose concentrations.

